# S-RNase Evolution in Self-Incompatibility: Phylogenomic Insights into Synteny with S-Like Genes

**DOI:** 10.1101/2024.08.14.607955

**Authors:** Yunxiao Liu, Yangxin Zhang, Songxue Han, Bocheng Guo, Jiakai Liang, Ze Yu, Fan Yang, Yaqiang Sun, Jiayu Xue, Zongcheng Lin, M. Eric Schranz, Changfei Guan, Fengwang Ma, Tao Zhao

## Abstract

S-RNases are essential in the gametophytic self-incompatibility (GSI) system of many flowering plants, where they act as stylar-S determinants. Despite their significance, the syntenic genomic origin and evolutionary trajectory of S-RNase genes in eudicots have remained largely unclear. Here, we performed large-scale phylogenetic and microsynteny network analyses of RNase T2 genes across 130 angiosperm genomes, encompassing 35 orders and 56 families. Based on the connections observed in the synteny network, particularly the persisting synteny block in none-SI Cucurbitaceae species, we infer that well-characterized S-RNase genes (Class III-A RNase genes) were duplicated and evolved from Class I S-like RNase genes, possibly as a result of the gamma triplication event shared by core eudicots. Additionally, we identified frequent lineage-specific gene transpositions of S-RNase genes across diverse angiosperm lineages, including Rosaceae, Solanaceae, and Rutaceae families, accompanied by a significant increase in transposable element (TE) activity near these genes. Our findings delineate the precise genomic origin and evolutionary path of eudicot S-RNase genes, enhancing our understanding of the evolution of the S-RNase-based GSI system.

## Introduction

Ribonucleases (RNases) are integral to a variety of cellular processes, including DNA replication, RNA metabolism, plant defense, and self-incompatibility (SI) (Luhtala & Parker, 2010; MacIntosh, 2011). The RNase T2 family, characterized by two conserved active sites, is particularly widespread across organisms and plays essential biological roles (Irie, 1999). In plants, the RNase T2 gene family has expanded significantly, with gene duplication and loss leading to varying numbers among species (MacIntosh et al., 2010). This family is divided into two subfamilies based on their role in gametophytic self-incompatibility (GSI) responses in plants: S-like RNases and S-RNases (MacIntosh et al., 2010). Phylogenetic analyses and intron count categorize them into three distinct groups (Igic and Kohn, 2001), where S-like RNases are assigned to Class I and II, while S-RNase genes belong to Class III. Despite structural similarities, they exhibit notable differences in gene expressions and functions. While S-like RNases are involved in gene expression control and antimicrobial defense responses, S-RNases are key determinants in the GSI systems of numerous flowering plants (angiosperms) lineages (Asquini et al., 2011; Franklin-Tong & Franklin, 2003; Hua et al., 2008; Liang et al., 2020; Ramanauskas & Igić, 2021; Takayama & Isogai, 2005).

The GSI mechanism serves as a widespread system that prevents inbreeding and promote outcrosses in bisexual flowering plant species (De Nettancourt, 2001). Current knowledge identifies four types of SI systems (Zhao et al., 2022). Type I GSI, observed in various plant families like Rosaceae, Solanaceae, Plantaginaceae, Rutaceae, Rubiaceae, and Cactaceae families, operates through S-RNase-based mechanisms. Type II SI, referred to as Sporophytic Self-Incompatibility (SSI), exists in Brassicaceae and relies on SRK and SCR proteins (Schopfer et al., 1999; Suzuki et al., 1999). Type III GSI, found in *Papaver*, is governed by PrsS and PrpS proteins (Foote et al., 1994; Wheeler et al., 2009). Type IV SI characterizes the sporophytic heterostyly of Primula (Giacomo et al., 2022; Huu et al., 2020). S-RNase-based GSI system is prevalent in eudicot species, as listed above, where it is controlled by the *S*-locus, which includes *S-RNase* genes as female determinants and *SLF* (*S*-locus F-box)/*SFB* (*S*-haplotype specific F-box) genes as male determinants, respectively, during SI interactions (Takayama & Isogai, 2005). The co-evolution of *S-RNase* and *SLF*/*SFB* genes is central to the GSI system, which is thought to have evolved once in eudicots (Kubo et al., 2010; Steinbachs & Holsinger, 2002; Vieira et al., 2008).

Extensive exploration of the RNase T2 gene, encompassing S-RNase genes, has been conducted across diverse plant species, including representatives of Poaceae and *Arabidopsis*, Rutaceae, Fabaceae, Rosaceae, and Plantaginaceae (Azizkhani et al., 2021; Igic and Kohn, 2001; Liang et al., 2017; MacIntosh et al., 2010; Morimoto et al., 2015; Vieira et al., 2021; Zhu et al., 2023). A considerable number of studies have notably emphasized the origin and functional divergence of S-RNase from T2-type RNases. These investigations employ phylogenomic approaches that leverage available genomic resources, compare genomic synteny, and infer phylogenetic relationships (Lv et al., 2022; Vieira et al., 2008; Zhao et al., 2022).

While the recent evolution of S-RNase genes of self-incompatibility systems in eudicots is widely acknowledged, the precise evolutionary processes leading to the current phylogenetic distribution of S-RNases from S like RNases remain elusive. Synteny information holds crucial importance in comparative genomic research, providing insights into gene origins. A microsynteny network, complementing phylogenetic analyses, reveals profound positional relationships between subfamilies at the genomic structure level (Zhao & Schranz, 2017; Zhao & Schranz, 2019). Here, we leverage available genomic data from 130 angiosperm species to conduct a comprehensive analysis of the RNase T2 gene family. Our aim is to elucidate the genomic origins and lineage-specific transpositions of S-RNases, offering novel insights into the intricate genomic trajectories and evolutionary history of this gene family.

## Results

### Phylogenetic analysis of the RNase T2 gene family in angiosperms

In our study, we analyzed the RNase T2 gene family across 130 fully sequenced plant species, representing diverse lineages and groups of flowering plants (Supplementary Fig. S1, Supplementary Table S1). Our analysis identified a total of 1366 RNase T2 genes (Supplementary Table S2). The maximum likelihood phylogenetic tree was reconstructed using the protein sequences of all identified RNase T2 genes, as well as the protein sequences of the S-RNase genes with reported functions (Supplementary Table S3). This tree classified these genes into three main clades: Class I (S-like RNase), Class II (S-like RNase), and Class III (including S-RNase), with seven major subclasses (Supplementary Fig. S2). This classification is consistent with current phylogenetic understanding (Igic and Kohn, 2001).

The three classes of RNase T2 genes showed distinct evolutionary patterns across angiosperm lineages. Class I genes were found in all major angiosperm groups, including eudicots, magnoliids, monocots, and the basal ANA grade, suggesting their preservation throughout angiosperm evolution (Supplementary Fig. S2). Class II genes were present in core angiosperms, except for the ANA grade. In contrast, Class III genes, including S-RNases, were exclusive to eudicots, with some lineages showing gene absence or loss, such as Brassicaceae and Asteraceae (Supplementary Fig. S2).

The RNase T2 gene family copy numbers varied significantly among angiosperm taxa, even within the same family, ranging from 3.3 copies in basal angiosperms to 12.9 copies in superasterids (Supplementary Fig. S2). Notably, some species like Durango root and cotton contained a higher number of genes, while others like jujube and begonia had only three genes each. (Supplementary Fig. S2).

The distribution of gene types within the RNase T2 family revealed a higher number of Class I (677 genes) and Class III (470 genes) genes compared to Class II (219 genes), indicating a smaller representation of Class II genes and a more significant expansion of Class I genes during plant evolution (Supplementary Fig. S2). Our observations indicate a moderate positive correlation between the gene numbers of Class I and Class II (Supplementary Fig. S3). However, we found weaker correlations between Class I and Class III genes as well as between Class II and Class III genes (Supplementary Fig. S3). We further classified this tree into seven major subclasses: Class I-A, Class I-B, Class II, Class III-A (which includes the reported S-RNase genes), Class III-B, Class III-C, and Class III-D, based on the phylogenetic relationships. The Class III-A represents the functional S-RNase subclade, the remaining subclades (Class III-B, Class III-C, and Class III-D) are non-functional S-like RNase subclades (Fig. 1, Supplementary Fig. S4).

**Figure 1.**
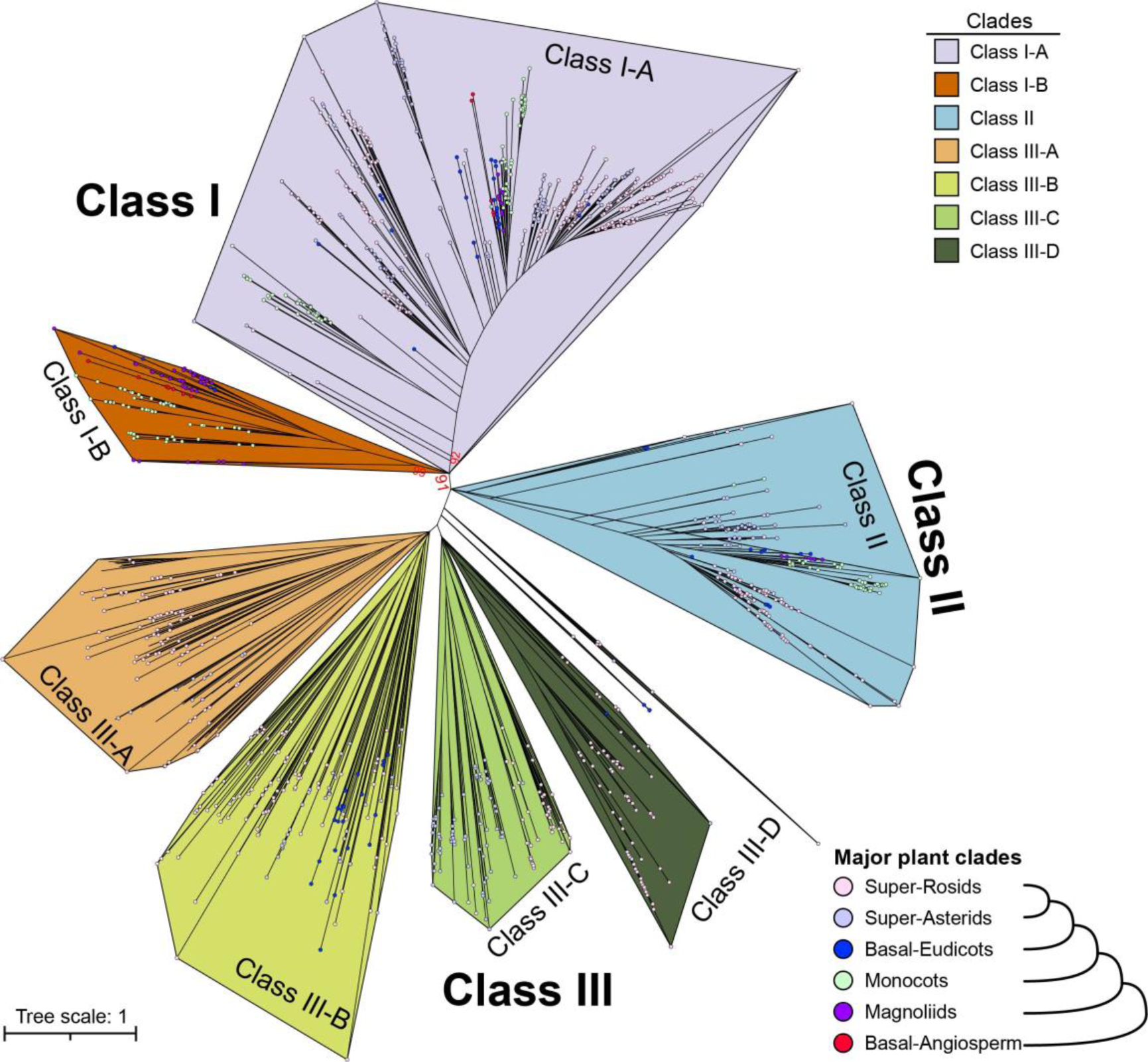
Phylogenetic relationships and distribution of the three major classes of the RNase T2 gene family in 130 angiosperm species. Phylogenetic tree reconstructed using the maximum likelihood (ML) method with 1,000 bootstrap replications based on full-length protein sequences of the RNase T2 gene family. The three major classes (seven subclasses): Class I, Class II, and Class III are distinguished. Notably, characterized S-RNase sequences from five families (Rosaceae, Solanaceae, Plantaginaceae, Rutaceae, and Rubiaceae) exhibiting gametophytic self-incompatibility (GSI) are clustered in Class III-A branch without colored dots. Colored dots denote different plant lineages: superrosids (light-pink), superasterids (purple), basal-eudicots (blue), monocots (light-green), magnoliids (purple), and basal-angiosperm (red). The ML tree displays bootstrap values for key nodes.

Five types of gene duplication events were identified in the RNase T2 gene family across 130 species (Supplementary Fig. S5). Different modes of duplication contributed to the amplification of specific gene classes in various plant lineages. Whole Genome Duplication (WGD) and Tandem Duplication (TD) events were major drivers for Class I gene expansion, particularly in Brassicaceae, Cleomaceae, and *Gossypium*. Transposed duplications (TRD) primarily contributed to the expansion of Class II genes in families like Rosaceae and Fabaceae. Dispersed duplications (DSD) were the main force in the evolution of Class III genes, including S-RNases (Supplementary Fig. S5, Supplementary Tables S4-5).

### Conservation of genomic contexts reveals shared ancestry between Class III S-RNase genes in Cucurbitaceae and Class I S-like RNase genes

We extracted the RNase T2 sub-network from the phylogenomic microsynteny database using all candidate RNase T2 genes. The network consisted of 793 nodes (RNase T2 genes) and 19873 edges (syntenic relationships) (Supplementary Fig. S6, Supplementary Table S6). Using the infomap cluster algorithm, we identified 67 synteny clusters. Class I genes showed more interconnections compared to other classes, suggesting better preservation of syntenic relationships. In contrast, Class III genes formed most small-sized clusters, indicating their active evolution of genomic contexts (Supplementary Fig. S7A). Gene syntenic retention rate (number of genes in synteny network relative to genes from the phylogenetic clade) and average clustering coefficient for Class III is the lowest (Supplementary Fig. S7B-E).

Mapping syntenic relationships onto the phylogenetic tree showed that, in general, synteny correlated well with phylogenetic classifications (Fig. 2A). Notably, we observed significant syntenic conservation between some Class III-A genes and Class I S-like RNase genes (Fig. 2A). Delving into the details of the relevant synteny clusters, we identified specific synteny between Class III-A genes in Cucurbitaceae, such as those found in *Cucumis sativus*, *Citrullus lanatus*, and *Cucurbita maxima*, and Class I genes from other angiosperm species (Fig. 2B, Supplementary Fig. S8A).

**Figure 2.**
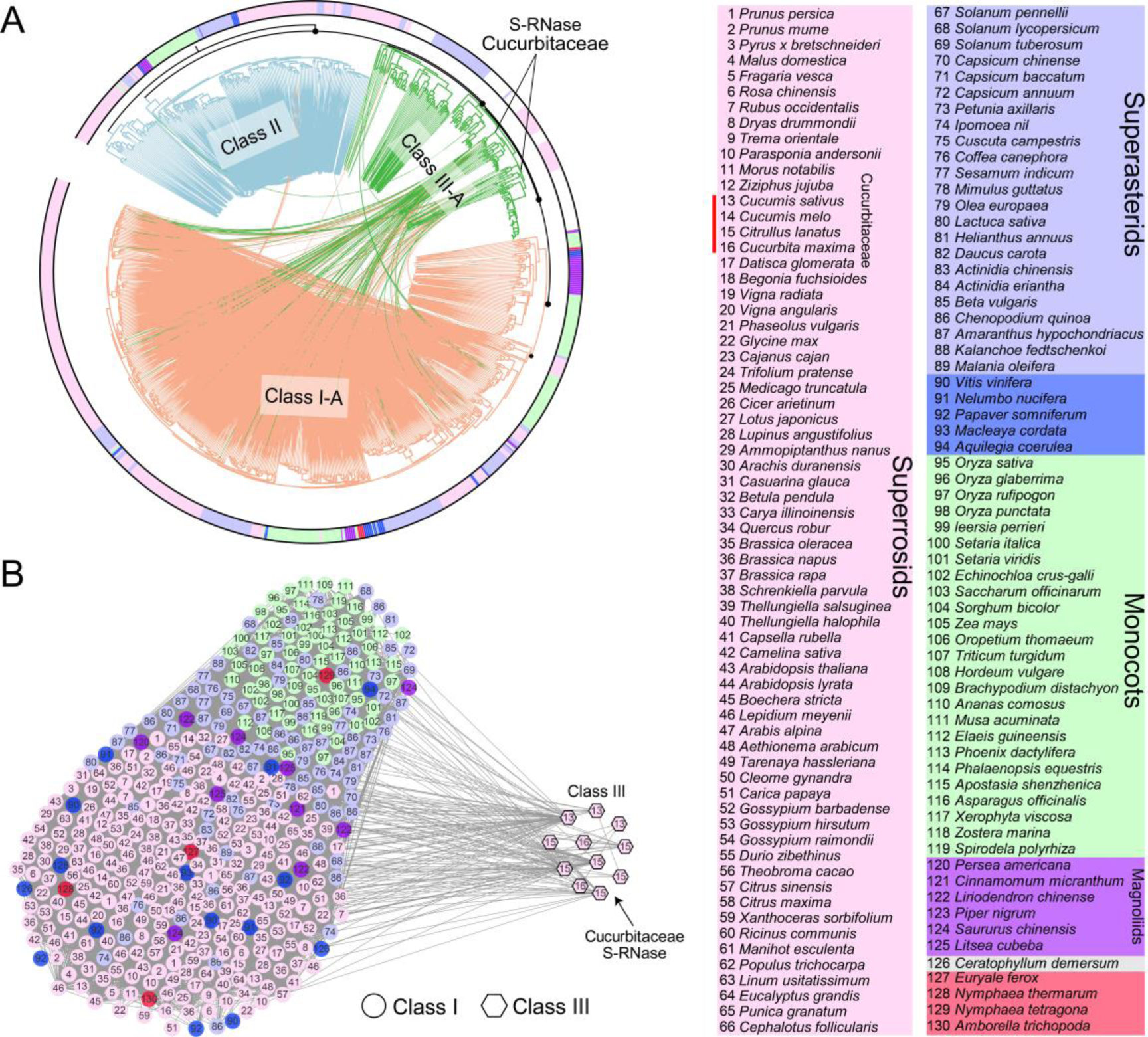
Phylogenomic synteny network analysis of the RNase T2 gene family in 130 angiosperm genomes. (**A**) The maximum-likelihood gene tree of the RNase T2 gene family accompanied by syntenic relationships between the genes. Genes devoid of connecting lines correspond to functional S-RNase sequences sourced from scientific literature. Notably, synteny relationships show links between Cucurbitaceae Class III genes and Class I genes. (**B**) Syntenic relationships for Class I and Class III RNase T2 genes within the phylogenomic context. The nodes are labeled and colored according to species and clades. Node shapes denote different phylogenetic clades or classes.

The phylogenetic tree of Class III genes showed that all function-reported S-RNase genes were predominantly clustered in Class III-A branch, indicating its representation of the S-RNase clade as mentioned above (Fig. 3A). Interestingly, Maleae S-RNase genes were closely related to Cucurbitaceae RNase T2 genes and received high overall bootstrap support values (BS = 99%) (Fig. 3B). We investigated the phylogenetic distributions of the Class III genes from all the 29 species with unisexual flowers in our dataset (Fig. 3C). The result showed that only 7 species contain genes from the Class III-A S-RNase clade (Fig. 3C), however, only the genes from Cucurbitaceae species can be found in synteny network.

**Figure 3.**
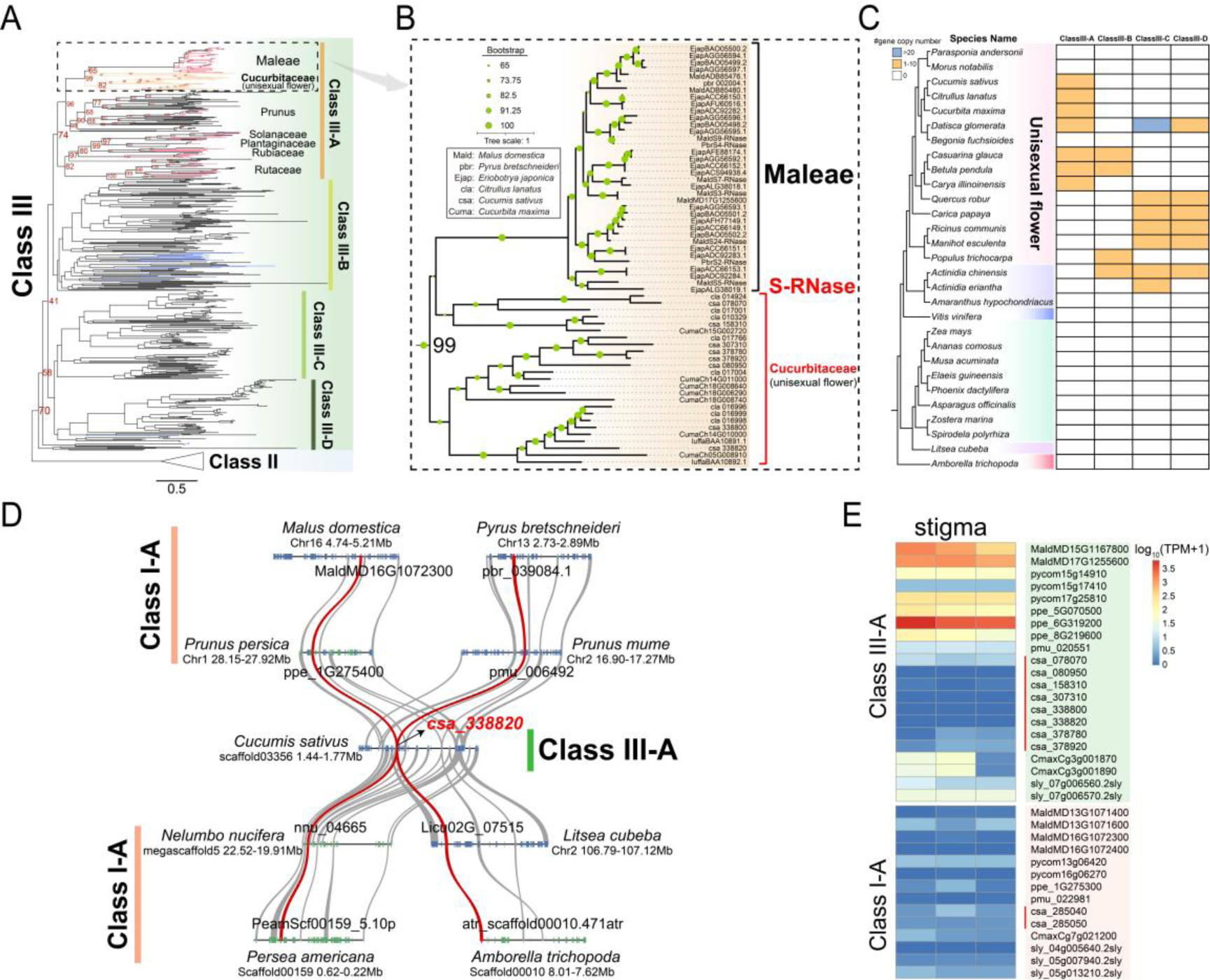
Synteny and phylogenetic analysis highlighting Class III S-RNase genes from Cucurbitaceae syntenic to Class I S-like RNase genes in angiosperms. (**A**) The maximum-likelihood (ML) tree illustrating Class III clades extracted from the comprehensive RNase T2 phylogeny depicted in Fig. 1. The clades representing functional S-RNase genes are highlighted in red, while clades representing Cucurbitaceae and basal-eudicots (e.g., *Aquilegia coerulea* and *Papaver somniferum*) are labeled in orange and blue, respectively. Ultrafast bootstrapping values are provided for key nodes. (**B**) An enlarged segment of the phylogenetic tree of the Class III genes from (**A**). Abbreviations of species names are displayed in the box at the upper left corner. (**C**) Copy number distribution of Class III subclasses among tested plant species with unisexual flowers. (**D**) Microsynteny relationships of *Cucumis sativus* S-RNase gene (csa_338820) and its syntenic genes in basal angiosperms (*Amborella trichopoda*), basal eudicots (*Nelumbo nucifera*), magnoliids (*Litsea cubeba* and *Persea americana*), and Rosaceae species (*Malus domestica, Pyrus bretschneideri, Prunus persica*, and *Prunus mume*). Curves linking the syntenic RNase T2 genes are highlighted in dark red. (**E**) Expression levels for RNase T2 genes in Class III-A and Class I-A in stigma.

We depicted some representative conserved synteny blocks between Class III-A S-RNase genes in Cucurbitaceae (e.g., cucumber csa_338820) and corresponding Class I genes from various species, including basal angiosperms and eudicots (Fig. 3D, Supplementary Fig. S8B). We then investigated the expression pattern of these Cucurbitaceae S-RNase genes and the syntenic Class I genes. Compared to the S-RNase genes from apple, pear, and peach, the non-functional orthologs of S-RNase genes in *Cucumis sativus* (e.g. csa_338820) exhibit low expression, which is similar to its syntenic Class I gene (csa_285040) (Fig. 3E). This suggests little functional divergence occurred between Cucurbitaceae Class III-A and Class I genes.

### The S-RNase genes of *Cucumis sativus* represent duplicates retained from the gamma triplication event

We exploit phylogenetic analysis and ancestral synteny block reconstruction to study the origin of Cucurbitaceae Class III-A S-RNase genes and Class I S-like RNase genes. Again, Cucurbitaceae RNase T2 genes can be classified into three Classes, while distinct collinearity relationships were found between Class III S-RNase genes and Class I RNase T2 genes, for example between the gene pairs of csa_338820 and csa_285040 (*Cucumis sativus*), and cla_016996 and cla_010187 (*Citrullus lanatus*) (Fig. 4A).

**Figure 4.**
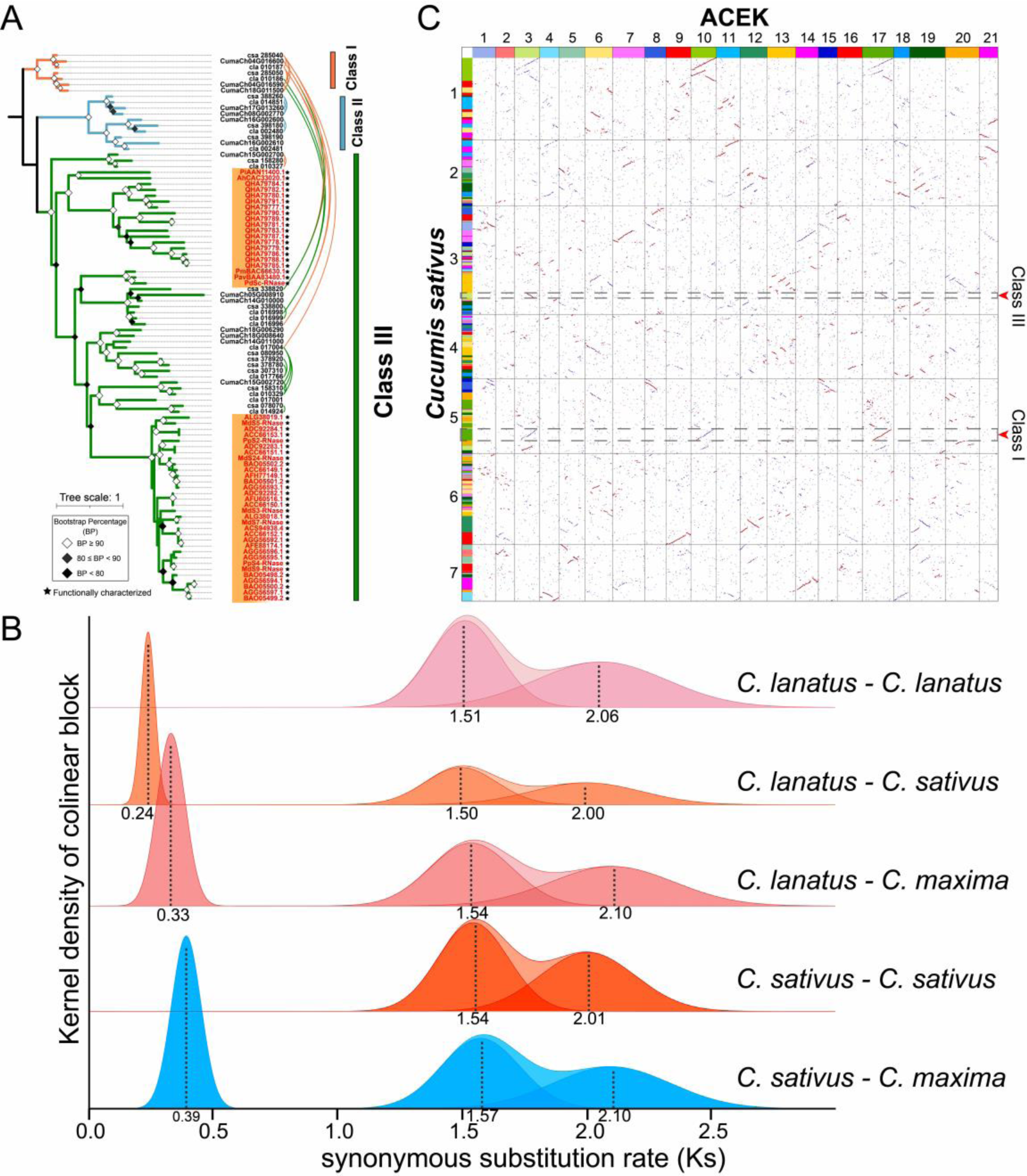
The Class III and Class I RNase T2 syntenic gene pairs in Cucurbitaceae species. (**A**) Maximum-likelihood gene tree of the RNase T2 gene family along with syntenic relationships among genes in Cucurbitaceae species studied. The phylogenetic tree is rooted using protein sequences of Class I genes as the outgroup. Genes marked in red labels and black stars indicate functional S-RNase genes. (**B**) Distributions of synonymous substitution rates (*Ks*) among collinear genes of the compared genomes. The curve is fitted using Gaussian mixture model, with peak values indicated in grey line. The *Ks* peaks around 1.5 correspond to the cucurbit-common tetraploidization event (CucWGD1), while the peaks around 2.0 correspond to the whole-genome triplication event (gamma) in core eudicots. (**C**) Homologous gene dot plots comparing the genome of *Cucumis sativus* with ACEK (ancestral core eudicots karyotype). Red dots represent the best BLAST-hits, while blue dots represent other BLAST-hits. The syntenic blocks of *Cucumis sativus* S-RNase gene (csa_338820) and Class I S-like gene (csa_285040) in ACEK are indicated by gray dashed box.

The synonymous nucleotide substitution rate (*Ks*) distributions of intragenomic syntenic gene pairs within tested Cucurbitaceae genomes of *Cucumis sativus*, *Citrullus lanatus*, and *Cucurbita maxima* revealed a consensus pattern of two polyploidization events (Fig. 4B). Specifically, the *Ks* ∼1.5 peaks correspond to the reported cucurbit-common tetraploidization event (CucWGD1) (Fig. 4B) (Ma et al., 2022), while the *Ks* ∼ 2.0 peaks caused by the whole genome triplication (gamma) event shared in core eudicots (Jiao et al., 2012). Considering the *K*s values of the ‘S-RNase-Class I’ syntenic pair we have pinpointed, namely 1.99 for csa_338820 and csa_285040, and 2.21 between cla_016996 and cla_010187, it is plausible to conclude that the origin of Cucurbitaceae S-RNase genes can be traced back to gamma triplication event.

Moreover, when mapping *Cucumis sativus* genome to Ancestral Core Eudicots Karyotype (ACEK), *Cucumis sativus* S-RNase gene csa_338820 (positioned on chromosome 3) and Class I S-like gene csa_285040 (positioned on chromosome 5) correspond to Chromosomes 3 and 17 of ACEK, respectively (Fig. 4C). This result again suggests their origin from the eudicot gamma triplication event.

### Noteworthy lineage-specific gene transpositions of S-RNase genes, which are accompanied by transposable element (TE) activities across angiosperms

We identified 67 synteny communities from the synteny network of the RNase T2 gene family. Class III genes (including S-RNases) from Rosaceae, Rutaceae, Solanaceae, and Malvaceae formed distinct synteny clusters specific to their family, as did Class II (S-like RNase) genes observed in Brassicaceae, Cucurbitaceae, and Poaceae (Fig. 5A, Supplementary Table S7).

**Figure 5.**
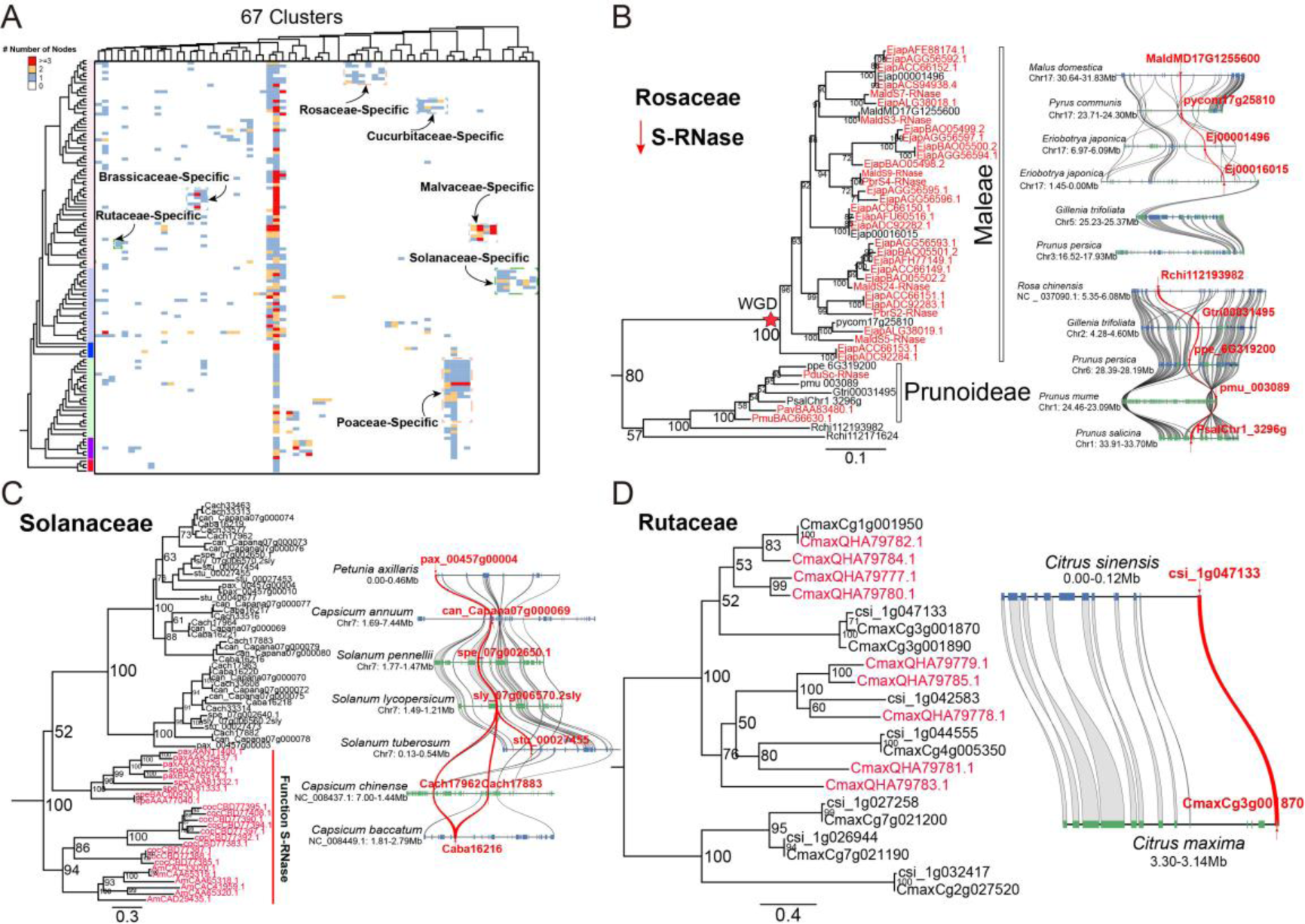
Insights into frequent gene transpositions of S-RNase genes in angiosperms. (**A**) Phylogenomic synteny profiling (copy-number profiling of microsynteny clusters across a phylogeny) of the synteny clusters of the RNase T2 gene family across 130 angiosperm genomes. Groups of lineage-specific clusters are boxed and labeled. The columns show synteny clusters and cell colors indicate the number of nodes (genes grouping in that cluster) per species. (**B-D**) Lineage-specific transpositions of Class III S-RNase genes in Rosaceae, Solanaceae, and Rutaceae, respectively. Reported functional S-RNase genes are highlighted in red, with gray curves connecting syntenic flanking genes. Genes located on the reverse strand are shown in green and genes located on the forward strand are shown in blue. S-RNase syntelogs are marked in red and connected by red lines.

As an example, a detailed synteny network analysis was performed for the RNase T2 genes in 11 representative Rosaceae species. The synteny network is comprised of 138 nodes and 239 edges, and 12 clusters were identified (Supplementary Tables S8-10). Phylogenetic and synteny analysis was performed to elucidate their evolutionary relationships (Fig. 5B, Supplementary Fig. S9-S10). Of note, S-RNase genes from Maleae species (*Malus domestica*, *Pyrus communis*, and *Eriobotrya japonica*), comparing with other Rosaceae species (*Gillenia trifoliata*, Prunoideae, and Rosoideae species), cluster in a specific synteny context and orthologous group (Fig. 5B, Supplementary Fig. S11-S13, Supplementary Tables S11-12). Besides Rosaceae, we also find such lineage-specific gene transpositions of S-RNase genes in Solanaceae (Fig. 5C) and Rutaceae species (Fig. 5D) as well. These findings suggest a frequent gene transposition of S-RNase genes in eudicot lineages. Upon closer inspection, long terminal repeat (LTR) retrotransposons were found adjacent to S-RNase genes in several plant genomes, including *Malus domestica*, *Solanum lycopersicum*, *Antirrhinum majus L*., and *Coffea canephora* (Supplementary Fig. S14, Supplementary Table S13). These findings highlight the impact of transposable elements to lineage-specific transposition of S-RNase genes, as well as the decreased synteny for Class III genes.

## Discussion

### Conserved synteny found between S-RNase and S-like RNase genes

Phylogenomic synteny analysis is becoming a crucial tool for elucidating the conserved gene order across multiple genomes, inferring functional relationships, and investigating the evolutionary lineage of genes (Ruelens et al., 2013; Zhao et al., 2017; Schultz et al., 2023). Despite the abundance of research on Class III S-RNase genes in certain plant families such as Rosaceae, Solanaceae, and Rutaceae, our study uncovered conserved syntenic relationships between Class III S-RNase and Class I S-like RNase genes. The obtained results have considerably enriched our understanding of complex evolutionary trajectories, shedding light on the existence of genomic ’fossils’, or remnants from previous evolutionary stages found within the well-conserved synteny in Cucurbitaceae species genomes.

Most species within the plant kingdom traditionally exhibit bisexual flowers (Matthews & Endress, 2004). Yet, the evolution of different mechanisms aimed at curtailing self-fertilization further corroborates the inherent tendency of flowering plants to promote cross-pollination. This promotes increased genetic diversity and the fitness of their offspring. The Cucurbitaceae family, comprising of important vegetables and melons, exhibits unisexuality across most of its member species, indicating a potential link between S-RNase genes and the loss of self-incompatibility (Boualem et al., 2015). It seems likely that Cucurbitaceae species have evolved to prevent self-pollination, exhibiting mechanisms such as monoecy, protandry, and dioecy (Steinbachs & Holsinger, 2002). As a result, the S-RNase-based GSI mechanism, prevalent in certain plant families, probably carries less relevance (Boualem et al., 2015).

Within this evolutionary context, the well-preserved conservation of genomic context (synteny) within the Cucurbitaceae species offers compelling evidence that the Class III genes, inclusive of S-RNase genes, have evolved from Class I genes. The study thereby underscores the presence of unisexual flowers and the absence of an S-RNase-based GSI mechanism, or similar reproductive systems, as likely contributing factors to such genomic conservation. These findings contribute valuable insights to our understanding of gene evolution mechanisms and plant reproductive system adaptations.

### Impact of transposable elements to S-RNase evolution

Dispersed duplication mode contributes most to Class III genes, which indicate their frequent translocations. Moreover, we identified lineage-specific clusters of Class III S-RNase genes in families like Rosaceae, Solanaceae, and Rutaceae, hinting at notable lineage-dependent transposition events of S-RNase genes. The Maleae-specific gene cluster seemingly aligns with the most recent and species-specific whole-genome duplication (WGD) event in Rosaceae (Velasco et al., 2010; Xiang et al., 2017).

TEs have often been implicated in inducing dispersed genes and transpositions, thus influencing gene composition, function, and as a result, the evolution of plant genomes (Lisch, 2013). The insertion of a MITE transposon near the S-RNase promoter resulted in the loss of SI traits in Rutaceae lineage (Hu et al., 2024). The proximity of LTR transposons to the S-RNase genes, particularly in the Rosaceae lineage, emphasizes the potential of these TEs to facilitate gene rearrangements and the transposition of S-RNase genes, either by introducing them to fresh genomic landscapes or by displacing them from their initial positions, thus contributing to the formation of two lineage-specific genomic contexts in Maleae and Prunoideae.

Interestingly, the diversity in operational mechanisms within the gametophytic self-incompatibility (GSI) system, marked by differential pollen recognition in two distinct branches of Rosaceae (Maleae and Prunoideae), appears to ride on the divergence in syntenic genomic contexts (Akagi et al., 2016; Ashkani & Rees, 2016; Fujii et al., 2016; Matsumoto & Tao, 2016). The significant genomic disparities introduced by TE insertion within S-RNase genes likely contribute to the mechanistic diversity observed within pollen recognition processes.

In conclusion, our study encapsulated a phylogenomic approach, integrated with an intensive synteny-based analysis, to shed light on the evolutionary trajectory of S-RNase genes. We unravelled the origins of Class III-A S-RNase genes in Cucurbitaceae from Class I S-like RNase genes and illuminated the specific genomic landscapes nurturing Class III S-RNase genes across different angiosperm lineages. This comprehensive understanding pertaining to the evolution of RNase T2 genes furthers our knowledge within the broader domains of plant reproductive biology and evolutionary genetics.

## Summary

Within this study, we proposed a model for the origins and evolutionary trajectory of S-RNase genes, drawing from the observations discussed above (Fig. 6). Previous research has demonstrated that Class I RNase T2 genes spanned all major land plant lineages, deriving initially from bryophytes, whereas Class II RNase T2 genes stemmed from the seed plants (Ramanauskas & Igić., 2017).

**Figure 6.**
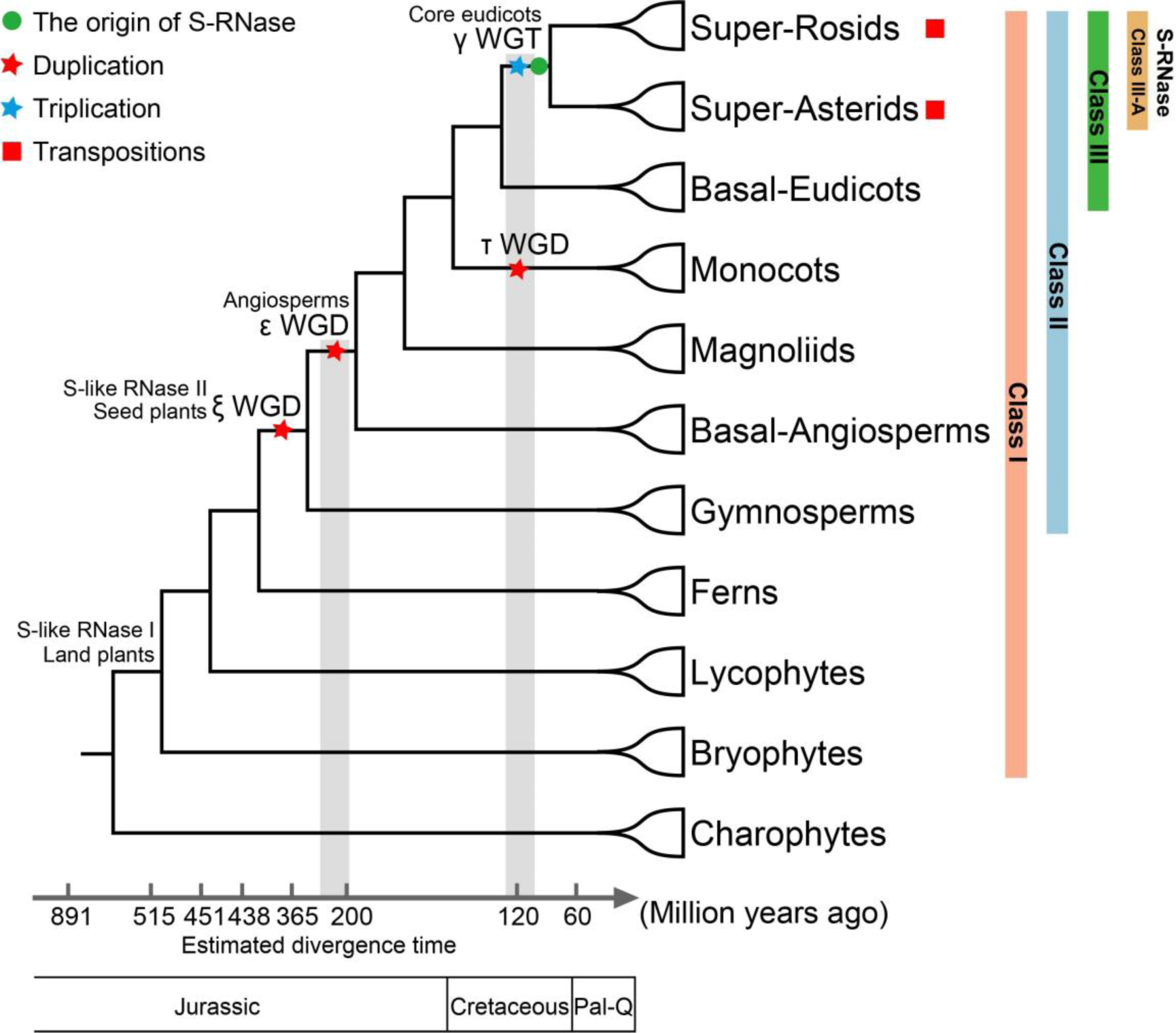
Evolution Trajectory of S-RNase genes in land plants. The figure outlines the hypothesized origin and diversification of S-RNase genes within the context of land plant evolution. The green circle represents the proposed origination of S-RNase genes, while the red squares denote instances of gene transpositions. Key whole-genome duplication (WGD) events are marked with red pentagrams, and whole-genome triplication (WGT) events with blue pentagrams, placed accordingly along the branches of the phylogenetic tree. The tree, depicted on the left, illustrates the evolutionary relationships among different plant clades. The timeline at the bottom provides estimated divergence times, calibrated according to the findings (Morris et al., 2018; Wu et al., 2020). The phylogenetic position of Magnoliids here corresponding to APG IV (Chase et al., 2016).

In contrast, the existence of Class III genes, encompassing the S-RNase genes, appears to be restricted to eudicots. Proceeding from this, a significant gamma triplication event, estimated to be around 120 million years old, set the precedent for the emergence of Class III-A (a Class III subclade) functional S-RNase genes from Class I S-like RNase genes. This emergence was corroborated by identified conserved microsynteny relationships. Following this, we noted frequent lineage-specific transpositions of S-RNase genes across diverse families.

Investigating the provenance of the S-RNase genes in *Cucumis sativus*, we identified these as duplicates maintained from the core eudicots gamma triplication event. A previous study alluded to this gamma (γ) event as restricted solely to core eudicots (Jiao et al., 2012). Along with this event, gamma paralogs of RNase T2 genes for basal eudicots underwent triple duplication, diversifying across three classes (Class I, Class II, and Class III). Intriguingly, in basal eudicots such as *Nelumbo nucifera*, *Papaver somniferum*, and *Aquilegia coerulea*, each of the three classes contains a copy of RNase T2 genes.

To the best of our knowledge, our study presents a compelling body of evidence, underpinned by genomic synteny, postulating that eudicot S-RNase genes evolved and multiplied from Class I S-like RNase genes, likely traceable back to the gamma palaeohexaploidy event. However, unraveling the precise timelines and mechanisms facilitating this extensive duplication demands further research and vigorous testing. Exploring the possible neofunctionalization or subfunctionalization of Class III genes derived from Class I genes in subsequent scenarios also emerges as an area of crucial importance.

## Materials and methods

### Plant genome resources and identification of RNase T2 Family genes

In our study, we analyzed 130 fully sequenced and annotated plant reference genomes across various taxa: 66 superrosids, 23 superasterids, 5 basal eudicots, 25 monocots, 6 Magnoliids, and 4 basal angiosperms. Within the basal angiosperms, predating the monocot-eudicot divergence, we incorporated *Amborella trichopoda*, *Euryale ferox*, *Nymphaea colorata*, and *Nymphaea thermarum* (Supplementary Fig. S1).

Genome annotations and coding sequences were sourced from established repositories like NCBI, Ensemble, GigaDB, CoGe, Phytozome, and specific databases such as GDR, Sol Genomics Network, Citrus Genome Database, and others (refer to Supplementary Table S1 for detailed links).

For broad and representative coverage, we compiled annotated protein sequences and genome annotation files with gene positional data from each genome. For coherent labeling in phylogenetic trees, species names were abbreviated into three or four-letter codes and paired with protein IDs. A list with genomes and corresponding specifics is available in Supplementary Table S1.

We have identified RNase T2 family genes using the seed alignment file of the RNase T2 domain (PF00445) from the Pfam database to form a Hidden Markov Model (HMM) file. Potential candidates were initially identified via HMMER3.0 (Finn et al., 2011), adopting the default inclusion threshold (E-value < 1E-3). The completeness and presence of the typical RNase T2 domain within protein sequences for candidate genes were further scrutinized (refer to http://www.smart.embl-heidelberg.de/ and http://pfam.xfam.org/). For a thorough list of the RNase T2 gene family members for 130 angiosperms genomes, see Supplementary Tables S2.

### Phylogenetic reconstruction

We obtained amino acid sequences of documented S-RNase and S-like RNase genes from GenBank (https://www.ncbi.nlm.nih.gov/genbank/) and compiled them in Supplementary Table S3. For the large-scale phylogenetic gene tree of the RNase T2 gene family across 130 angiosperm species, full-length amino acid sequences were aligned with MAFFT (version 7.475), using the (-localpair –maxiterate 1000 --thread 20 –reorder) parameters and the L-INS-I strategy (Katoh & Standley, 2013). This alignment was subsequently manually curated in MEGA (version 11) to eliminate gaps and discard sequences without conserved motifs (Tamura et al., 2011). Using the ModelFinder algorithm in IQ-TREE, optimal amino acid substitution models were determined via the approximate maximum likelihood method (-m MF -T AUTO) where the ’WAG+R8’ model was identified as the best-fit model for the RNase T2 gene family across 130 species. (Kalyaanamoorthy et al., 2017; Nguyen et al., 2015). The maximum likelihood (ML) phylogenetic tree was then computed with IQ-TREE (version 2.0.3) applying the parameters: -m WAG+R8 -alrt 1000 -bb 1000 -nt 10 (Lin et al., 2013). Additionally, for the small-scale phylogenetic tree of 11 Rosaceae species, the result was inferred with IQ-TREE (version 2.0.3) using the parameters: -m VT+R6 -alrt 1000 -bb 1000 -nt 10. The resulting maximum-likelihood gene trees were subsequently visualized and annotated with the online tools iTOL v.5 (https://itol.embl.de/) and Figtree v1.4.4 (http://tree.bio.ed.ac.uk/software/figtree/) (Letunic & Bork, 2021).

### Determination of gene duplication modes

The origins of RNase T2 genes in each species were categorized into five different modes of gene duplication for this study: whole-genome duplication (WGD), tandem duplication (TD), proximal duplication (PD), transposed duplication (TRD), and dispersed duplication (DSD). This categorization utilized DupGen_finder (https://github.com/qiao-xin/DupGen_finder) with default parameters (Qiao et al., 2019).

Specifically, tandem duplicates (TD) refer to gene copies located immediately adjacent to each other within 10 annotated genes on the same chromosome, devoid of any interspersed paralogs. Proximal gene pairs (PD) are non-tandem pairs separated by 10 or fewer genes on the same chromosome. Transposed duplication (TRD), often labeled as ’transposed,’ entails distally located pairs where one gene is syntenic and the other is non-syntenic. This arrangement creates a gene pair comprising an ancestral and a novel locus. Dispersed duplication (DSD) events generate two gene copies which are neither adjacent nor colinear.

Any remaining duplicates not falling under WGD, tandem, proximal, and transposed duplicates were considered as dispersed duplicates. The precedence of duplicated genes was established as follows: WGD>TD>PD>TRD>DSD. Singletons denote genes within a species lacking homologous counterparts in the target species and are not classified as a type of duplication. Supplementary Tables S4-5 contain the results of this categorization.

### Synteny network analysis and phylogenomic profiling of synteny clusters

The SynNet construction pipeline (https://github.com/zhaotao1987/SynNet-Pipeline), developed by Zhao and Schranz (2017), was employed to build synteny networks, which encompass intra- and inter-genome synteny comparisons and synteny relationships among all the genes from the studied angiosperm genomes. The first step involved an all-against-all reciprocal comparison of protein sequences from each of the 130 angiosperms species under study using Diamond v3.3.2 (Buchfink et al., 2015). Next, we used MCScanX to calculate genomic collinearity (i.e. conserved gene order and content across multiple species genomes) between all pairwise genome combinations under default parameters (minimum match size for a collinear block = 5 genes, max gaps allowed = 25 genes) (Wang et al., 2012).

Following that, with the gene list of all candidate RNase T2 genes, a sub-network specifically for the RNase T2 genes was extracted from the entire synteny network database (Supplementary Table S6). Each node in this sub-network represents a gene, edges signify syntenic connections between genes, and edge lengths are disregarded (unweighted). To cluster the network, we implemented the infomap algorithm (map equation framework) (Rosvall & Bergstrom, 2008).

The R package "igraph" was used to perform network statistical analysis. The resulting RNase T2 gene family synteny network was visualized and analyzed using Cytoscape v.3.8.0 (Shannon et al., 2003) and Gephi (Bastian et al., 2009).

We used the iTOL v.5 web server (https://itol.embl.de/) to map syntenic relations onto the constructed phylogenetic tree, and prepared phylogenomic profiles for each synteny cluster, recording the number of nodes for each species per cluster. We assessed the copy number of syntenic genes across species within each community. These profiles, illustrating the count of syntenic RNase T2 genes in each genome for each synteny cluster, were assembled and visualized using "Jaccard" distance and ’ward.D’ clustering. More extensive information on the synteny network analysis and resulting profiles are provided in Supplementary Tables S7-10.

We then displayed notable microsynteny contexts particularly focusing on the S-RNase genes among chosen species using MCscan (Python version) implemented in the jcvi package (https://github.com/tanghaibao/jcvi/wiki/MCscan-(Python-version)), featuring a window size of 40 genes to represent the flanking areas of the RNase T2 genes (Tang et al., 2015). Further details can be found in Supplementary Table S11.

### Ortholog identification and phylogenomic profiling of orthologous clusters

The Rosaceae dataset in this study consisted of eleven genomes from various Rosaceae species, showcasing three subfamilies: Amygdaloideae (*Malus domestica*, *Pyrus communis*, *Eriobotrya japonica*, *Gillenia trifoliata*, *Prunus persica*, *Prunus mume*, and *Prunus salicina*), Rosoideae (*Rubus occidentalis*, *Rosa chinensis*, and *Fragaria vesca*), and Dryadoideae (*Dryas drummondii*). We used OrthoFinder (version 2.5.2) with specific parameters (-S diamond -M msa -A mafft -I 1.5) to identify orthogroups among the protein sequences of these eleven Rosaceae species (Emms & Kelly, 2019).

From this analysis, we obtained a phylogenomic profile matrix comprising 28,069 non-redundant multigene clusters (orthogroups). Each orthogroup and each synteny cluster was annotated with the number of represented species to determine the dissimilarity among all the clusters. We then computed a dissimilarity index and executed hierarchical clustering using the "ward.D" method. The resulting cluster heatmap was visualized using "pheatmap", and detailed information about the identified orthogroups can be found in Supplementary Table S12.

### Synonymous substitution rates (*Ks*) calculation, identification of transposons

For the Cucurbitaceae genome, we produced *Ks* dot plots using the WGDI v.0.62 toolkit (Sun et al., 2022). We fashioned density distribution curves for *Ks* values using Kspeaks (-kp), and performed multipeak fitting with PeaksFit (-pf) software. We used the estimated mean peak values obtained from these curves to date Whole-Genome Duplication (WGD) events. For ACEK mapping, we applied the "-icl" parameter to recognize collinear genes between ACEK and specific species and the "-km" parameter to map ACEK. Finally, we utilized WGDI together with the "-d" parameter, adding the ancestor_left (*Cucumis sativus*) to produce the homologous dot plot.

We utilized the extensive de-novo TE annotator (EDTA version 1.9.4), with the parameter: --sensitive 1 --anno 1, to identify transposable elements (TEs) (Su et al., 2021). For our study, we only considered intact TEs. More detailed information about *Ks* values in the compared Cucurbitaceae genomes and the identified TEs is available in Supplementary Table S13.

### Gene expression analysis

The RNA-Seq gene expression data for this study was obtained from the public NCBI SRA database (https://www.ncbi.nlm.nih.gov/sra). We performed quality checks on the raw reads using Fastp v.0.12.4, applying parameters (-f 12 -F 12 -l 50) to ensure data cleanliness and high quality (Chen et al., 2018). The cleaned, high-quality reads derived from the samples were then mapped to the coding sequences (CDS) of the reference genomes.

Following this procedure, transcripts per million (TPM) gene expression values were computed using Kallisto v.0.46.2 and were log10-transformed for easier interpretation (Bray et al., 2016). The TPM values were used to quantify the expression levels of genes within tissues. Comprehensive details regarding the RNA-seq data used can be found in Supplementary Table S14.

### Accession numbers

Accession numbers of the sequence data from this article can be found in Supplemental Tables S1 (genome assemblies) and S2-3 (all RNase T2 genes).

## Acknowledgments

We thank High-Performance Computing (HPC) of Northwest A&F University (NWAFU) for providing computing resources.

## Author contributions

TZ, FM, CG, and YS designed the study. YL, YZ, HS, BG, LJ, ZY, and FY conducted the analyses, TZ, YL, JX, ZL, and MES analyzed data. YL and TZ wrote the paper. All co-authors read and edited the manuscript.

## Supplemental data

The following materials are available in the online version of this article.

**Supplemental Figure S1**. Phylogenetic relationships of the 130 angiosperm and 11 Rosaceae genomes analyzed in this study.

**Supplemental Figure S2**. Distribution of the three major classes of the RNase T2 gene family in 130 angiosperm species.

**Supplemental Figure S3**. Gene counts of different classes across different clades and correlation analysis.

**Supplemental Figure S4**. Phylogenetic relationships of the RNase T2 gene family in 130 angiosperm species.

**Supplemental Figure S5**. Modes of gene duplications for the RNase T2 gene family in 130 angiosperm species.

**Supplemental Figure S6**. A network view depicting all the syntenic relationships for the entire RNase T2 gene family from 130 angiosperm species.

**Supplemental Figure S7**. The network statistic for each class of the RNase T2 gene family.

**Supplemental Figure S8**. Synteny and phylogenetic analysis highlighting Class III S-RNase genes from Cucurbitaceae syntenic to Class I S-like genes in angiosperms.

**Supplemental Figure S9**. Phylogenetic tree of the RNase T2 gene family in Rosaceae.

**Supplemental Figure S10**. Microsynteny network analysis of the RNase T2 genes in Rosaceae genomes.

**Supplemental Figure S11**. Phylogenomic profiling of the microsynteny clusters from 11 Rosaceae species.

**Supplemental Figure S12**. Maximum-likelihood phylogenetic tree of Rosaceae RNase T2 genes and corresponding synteny network clusters.

**Supplemental Figure S13**. Phylogenomic profiling of the orthologous gene clusters of 11 Rosaceae species.

**Supplemental Figure S14**. Bar chart displaying the distribution of TEs around the S-RNase genes in the genomes of diverse eudicot species.

**Supplementary Table S1**. Comprehensive genome annotation details for the 130 angiosperm species used in this study.

**Supplementary Table S2**. Gene counts of the RNase T2 gene family members in each subclass and amino acid sequences of the identified RNase T2 genes.

**Supplementary Table S3**. Amino acid sequences of known S-RNase and S-like RNase genes.

**Supplementary Table S4**. Count of genes exhibiting various duplication modes within the RNase T2 gene family.

**Supplementary Table S5**. Count of genes categorized by various duplication modes within the RNase T2 gene family across different classes.

**Supplementary Table S6**. The full edgelist of the RNase T2 genes synteny network across 130 angiosperm species.

**Supplementary Table S7**. Phylogenomic profiling of the synteny clusters of the RNase T2 gene family.

**Supplementary Table S8.** RNase T2 genes identified in 11 Rosaceae species.

**Supplementary Table S9.** The edgelist of the RNase T2 synteny network of 11 Rosaceae species.

**Supplementary Table S10**. Phylogenomic profiling of the synteny clusters of the RNase T2 genes in 11 Rosaceae species.

**Supplementary Table S11**. Microsynteny relationships of the S-RNase genes in Maleae and *Prunus* species, referenced by the genome of *Malus domestica*.

**Supplementary Table S12.** Phylogenomic profiling of the orthogroups of the RNase T2 gene family in Rosaceae.

**Supplementary Table S13**. Information of *Ks* distributions among Cucurbitaceae genomes and the identified transposable elements (TEs) near S-RNases.

**Supplementary Table S14**. RNA-seq data used in this study.

## Data availability

All the data supporting the research presented in this study are included in the article or in the Supplemental data.

## References

Akagi, T., Henry, I. M., Morimoto, T., & Tao, R. (2016). Insights into the prunus-specific S-RNase-based self-incompatibility system from a genome-wide analysis of the evolutionary radiation of S Locus-related F-box genes. Plant Cell Physiol. 57(6), 1281–1294. 10.1093/pcp/pcw077.

Ashkani, J., & Rees, D. J. (2016). A comprehensive study of molecular evolution at the self-incompatibility locus of Rosaceae. J Mol Evol, 82(2-3), 128–145. 10.1007/s00239-015-9726-4.

Asquini, E., Gerdol, M., Gasperini, D., Igic, B., Graziosi, G., & Pallavicini, A. (2011). S-RNase-like sequences in styles of coffea (Rubiaceae). evidence for S-RNase based gametophytic self-incompatibility? Trop. Plant Biol. 4(3), 237–249. 10.1007/s12042-011-9085-2.

Azizkhani, N., Mirzaei, S., & Torkzadeh-Mahani, M. (2021). Genome-wide identification and characterization of legume T2 Ribonuclease gene family and analysis of GmaRNS9, a soybean T2 Ribonuclease gene, function in nodulation. 3 Biotech, **11**(12), 495. 10.1007/s13205-021-03025-x.

Bastian, M., Heymann, S., & Jacomy, M. (2009). Gephi: An open source software for exploring and manipulating networks. Proc. Int. AAAI Conf. Weblogs Soc. Media 8: 361–362. 10.13140/2.1.1341.1520.

Boualem, A., Troadec, C., Camps, C., Lemhemdi, A., Morin, H., Sari, M. A., Fraenkel-Zagouri, R., Kovalski, I., Dogimont, C., Perl-Treves, R., & Bendahmane, A. (2015). A cucurbit androecy gene reveals how unisexual flowers develop and dioecy emerges. Science, 350(6261), 688–691. 10.1126/science.aac8370.

Bray, N. L., Pimentel, H., Melsted, P., & Pachter, L. (2016). Near-optimal probabilistic RNA-seq quantification. Nat Biotechnol. 34(5), 525–527. 10.1038/nbt.3519.

Buchfink, B., Xie, C. & Huson, D. H. Fast and sensitive protein alignment using DIAMOND. Nat Methods. 12, 59–60 (2015). 10.1038/nmeth.3176.

Chase, M. W., Christenhusz, M. J. M., Fay, M. F., Byng, J. W., Judd, W. S., Soltis, D. E., Mabberley, D. J., Sennikov, A. N., Soltis, P. S., & Stevens, P. F. (2016). An update of the Angiosperm Phylogeny Group classification for the orders and families of flowering plants: APG IV. Bot. J. Linn. Soc, 181(1), 1–20. 10.1111/boj.12385.

Chen, S., Zhou, Y., Chen, Y., & Gu, J. (2018). fastp: an ultra-fast all-in-one FASTQ preprocessor. Bioinformatics, 34(17), i884–i890. 10.1093/bioinformatics/bty560.

De Nettancourt, D. (2001). Incompatibility and incongruity in wild and cultivated plants. Springer-Verlag, Berlin. 10.1007/978-3-662-04502-2.

Emms, D. M., & Kelly, S. (2019). OrthoFinder: phylogenetic orthology inference for comparative genomics. Genome Biol. 20(1), 238. 10.1186/s13059-019-1832-y.

Finn, R. D., Clements, J., & Eddy, S. R. (2011). HMMER web server: interactive sequence similarity searching. Nucleic Acids Res. 39(Web Server issue), W29-37. 10.1093/nar/gkr367.

Foote, H. C., Ride, J. P., Franklin-Tong, V. E., Walker, E. A., Lawrence, M. J., & Franklin, F. C. (1994). Cloning and expression of a distinctive class of self-incompatibility (S) gene from *Papaver rhoeas L*. Proc Natl Acad Sci U S A. 91(6), 2265–2269. 10.1073/pnas.91.6.2265.

Franklin-Tong, N. V., & Franklin, F. C. (2003). Gametophytic self-incompatibility inhibits pollen tube growth using different mechanisms. Trends in Plant Science, 8(12), 598–605. 0.1016/j.tplants.2003.10.008.

Fujii, S., Kubo, K., & Takayama, S. (2016). Non-self- and self-recognition models in plant self-incompatibility. Nat Plants. 2(9), 16130. 10.1038/nplants.2016.130.

Giacomo, P., Léveillé-Bourret, É., Yousefi, N., Choudhury, R. R., Keller, B., Diop, S. I., Duijsings, D., Pirovano, W., Lenhard, M., Szövényi, P., & Conti, E. (2022). Comparative genomics elucidates the origin of a supergene controlling floral heteromorphism. Mol Biol Evol. 39(2), msac035. 10.1093/molbev/msac035.

Hu, J., Liu, C., Du, Z., Guo, F., Song, D., Wang, N., Wei, Z., Jiang, J., Cao, Z., Shi, C., Zhang, S., Zhu, C., Chen, P., Larkin, R. M., Lin, Z., Xu, Q., Ye, J., Deng, X., Bosch, M., Franklin-Tong, V. E., & Chai, L. (2024). Transposable elements cause the loss of self-incompatibility in citrus. Plant Biotechnol J, 22(5), 1113–1131. 10.1111/pbi.14250.

Hua, Z. H., Fields, A., & Kao, T. H. (2008). Biochemical models for S-RNase-based self-incompatibility. Mol Plant. 1(4), 575–585. 10.1093/mp/ssn032.

Huu, C. N., Keller, B., Conti, E., Kappel, C., & Lenhard, M. (2020). Supergene evolution via stepwise duplications and neofunctionalization of a floral-organ identity gene. Proc Natl Acad Sci U S A. 117(37), 23148–23157. 10.1073/pnas.2006296117.

Igic, B., & Kohn, J. R. (2001). Evolutionary relationships among self-incompatibility RNases. Proc Natl Acad Sci U S A. 98(23), 13167–13171. 10.1073/pnas.231386798.

Irie, M. (1999). Structure-function relationships of acid ribonucleases: lysosomal, vacuolar, and periplasmic enzymes. Pharmacol Ther. 81(2), 77–89. 10.1016/s0163-7258(98)00035-7.

Jiao, Y., Leebens-Mack, J., Ayyampalayam, S., Bowers, J. E., McKain, M. R., McNeal, J., Rolf, M., Ruzicka, D. R., Wafula, E., Wickett, N. J., Wu, X., Zhang, Y., Wang, J., Zhang, Y., Carpenter, E. J., Deyholos, M. K., Kutchan, T. M., Chanderbali, A. S., Soltis, P. S., Stevenson, D. W., McCombie, R., Pires, J. C., Wong, G. K., Soltis, D. E., & Depamphilis, C. W. (2012). A genome triplication associated with early diversification of the core eudicots. Genome Biol. 13(1), R3. 10.1186/gb-2012-13-1-r3.

Kalyaanamoorthy, S., Minh, B. Q., Wong, T. K. F., von Haeseler, A., & Jermiin, L. S. (2017). ModelFinder: fast model selection for accurate phylogenetic estimates. Nat Methods. 14(6), 587–589. 10.1038/nmeth.4285.

Katoh, K., & Standley, D. M. (2013). MAFFT multiple sequence alignment software version 7: improvements in performance and usability. Mol Biol Evol. 30(4), 772–780. 10.1093/molbev/mst010.

Kubo, K., Entani, T., Takara, A., Wang, N., Fields, A. M., Hua, Z., Toyoda, M., Kawashima, S., Ando, T., Isogai, A., Kao, T. H., & Takayama, S. (2010). Collaborative non-self recognition system in S-RNase-based self-incompatibility. Science, 330(6005), 796–799. 10.1126/science.1195243.

Letunic, I., & Bork, P. (2021). Interactive Tree Of Life (iTOL) v5: an online tool for phylogenetic tree display and annotation. Nucleic Acids Res. 49(W1), W293–w296. 10.1093/nar/gkab301.

Liang, M., Cao, Z., Zhu, A., Liu, Y., Tao, M., Yang, H., Xu, Q., Jr., Wang, S., Liu, J., Li, Y., Chen, C., Xie, Z., Deng, C., Ye, J., Guo, W., Xu, Q., Xia, R., Larkin, R. M., Deng, X., Bosch, M., Franklin-Tong, V. E., & Chai, L. (2020). Evolution of self-compatibility by a mutant S(m)-RNase in citrus. Nat Plants. 6(2), 131–142. 10.1038/s41477-020-0597-3.

Liang, M., Yang, W., Su, S., Fu, L., Yi, H., Chen, C., Deng, X., & Chai, L. (2017). Genome-wide identification and functional analysis of S-RNase involved in the self-incompatibility of citrus. Mol Genet Genomics. 292(2), 325–341. 10.1007/s00438-016-1279-8.

Lin, Y., Hu, F., Tang, J., & Moret, B. M. (2013). Maximum likelihood phylogenetic reconstruction from high-resolution whole-genome data and a tree of 68 eukaryotes. Pac Symp Biocomput. 285–296. 10.1142/9789814447973_0028.

Lisch, D. (2013). How important are transposons for plant evolution? Nat. Rev. Genet, 14(1), 49–61. 10.1038/nrg3374.

Luhtala, N., & Parker, R. (2010). T2 Family ribonucleases: ancient enzymes with diverse roles. Trends Biochem Sci. 35(5), 253–259. 10.1016/j.tibs.2010.02.002.

Lv, S., Qiao, X., Zhang, W., Li, Q., Wang, P., Zhang, S., & Wu, J. (2022). The origin and evolution of RNase T2 Family and gametophytic self-incompatibility system in plants. Genome Biol Evol. 14(7). 10.1093/gbe/evac093.

MacIntosh. (2011). RNase T2 Family: Enzymatic Properties, Functional Diversity, and Evolution of Ancient Ribonucleases. Ribonucleases, 89–114. 10.1007/978-3-642-21078-5_4.

MacIntosh, Hillwig, M. S., Meyer, A., & Flagel, L. (2010). RNase T2 genes from rice and the evolution of secretory ribonucleases in plants. Mol Genet Genomics. 283(4), 381–396. 10.1007/s00438-010-0524-9.

Ma, L., Wang, Q., Zheng, Y., Guo, J., Yuan, S., Fu, A., Bai, C., Zhao, X., Zheng, S., Wen, C., Guo, S., Gao, L., Grierson, D., Zuo, J., & Xu, Y. (2022). Cucurbitaceae genome evolution, gene function and molecular breeding. Hortic Res. 9. 10.1093/hr/uhab057.

Matsumoto, D., & Tao, R. (2016). Distinct Self-recognition in the Prunus S-RNase-based Gametophytic Self-incompatibility System. Hort. J. 85, 289–305. 10.2503/hortj.MI-IR06.

Matthews ML., & Endress PK. (2004). Comparative floral structure and systematics in Cucurbitales (Corynocarpaceae, Coriariaceae, Tetramelaceae, Datiscaceae, Begoniaceae, Cucurbitaceae, Anisophylleaceae). Bot J Linn Soc. 145(2), 129–185. 10.1111/j.1095-8339.2003.00281.x.

Morimoto, T., Akagi, T., & Tao, R. (2015). Evolutionary analysis of genes for S-RNase-based self-incompatibility reveals S Locus duplications in the ancestral Rosaceae. Hort. J. 84. 10.2503/hortj.MI-060.

Morris, J. L., Puttick, M. N., Clark, J. W., Edwards, D., Kenrick, P., Pressel, S., Wellman, C. H., Yang, Z., Schneider, H., & Donoghue, P. C. J. (2018). The timescale of early land plant evolution. Proc Natl Acad Sci U S A, 115(10), E2274–e2283. 10.1073/pnas.1719588115.

Nguyen, L.-T., Schmidt, H. A., von Haeseler, A., & Minh, B. Q. (2015). IQ-TREE: A fast and effective stochastic algorithm for estimating maximum-likelihood phylogenies. Mol Biol Evol. 32(1), 268–274. 10.1093/molbev/msu300.

Qiao, X., Li, Q., Yin, H., Qi, K., Li, L., Wang, R., Zhang, S., & Paterson, A. H. (2019). Gene duplication and evolution in recurring polyploidization-diploidization cycles in plants. Genome Biol. 20(1), 38. 10.1186/s13059-019-1650-2.

Ramanauskas, K., & Igić, B. (2017). The evolutionary history of plant T2/S-type ribonucleases. PeerJ, 5, e3790. 10.7717/peerj.3790.

Ramanauskas, K., & Igić, B. (2021). RNase-based self-incompatibility in cacti. New Phytol. 231(5), 2039–2049. 10.1111/nph.17541.

Rosvall, M., & Bergstrom, C. T. (2008). Maps of random walks on complex networks reveal community structure. Proc Natl Acad Sci U S A. 105(4), 1118–1123. 10.1073/pnas.0706851105.

Ruelens, P., de Maagd, R. A., Proost, S., Theißen, G., Geuten, K., & Kaufmann, K. (2013). FLOWERING LOCUS C in monocots and the tandem origin of angiosperm-specific MADS-box genes. Nat Commun. 4, 2280. 10.1038/ncomms3280.

Schopfer, C. R., Nasrallah, M. E., & Nasrallah, J. B. (1999). The male determinant of self-incompatibility in Brassica. Science, 286(5445), 1697–1700. 10.1126/science.286.5445.1697.

Schultz, D. T., Haddock, S. H. D., Bredeson, J. V., Green, R. E., Simakov, O., & Rokhsar, D. S. (2023). Ancient gene linkages support ctenophores as sister to other animals. Nature, 618(7963), 110–117. 10.1038/s41586-023-05936-6.

Shannon, P., Markiel, A., Ozier, O., Baliga, N. S., Wang, J. T., Ramage, D., Amin, N., Schwikowski, B., & Ideker, T. (2003). Cytoscape: a software environment for integrated models of biomolecular interaction networks. Genome Res. 13(11), 2498–2504. 10.1101/gr.1239303.

Steinbachs, J. E., & Holsinger, K. E. (2002). S-RNase–mediated gametophytic self-Incompatibility is ancestral in eudicots. Mol Biol Evol. 19(6), 825–829. 10.1093/oxfordjournals.molbev.a004139.

Sun, P., Jiao, B., Yang, Y., Shan, L., Li, T., Li, X., Xi, Z., Wang, X., & Liu, J. (2022). WGDI: A user-friendly toolkit for evolutionary analyses of whole-genome duplications and ancestral karyotypes. Mol Plant. 15(12), 1841–1851. 10.1016/j.molp.2022.10.018.

Su, W., Ou, S., Hufford, M. B., & Peterson, T. (2021). A tutorial of EDTA: extensive De Novo TE annotator. Methods Mol Biol. 2250, 55–67. 10.1007/978-1-0716-1134-0_4.

Suzuki, G., Kai, N., Hirose, T., Fukui, K., Nishio, T., Takayama, S., Isogai, A., Watanabe, M., & Hinata, K. (1999). Genomic organization of the S locus: identification and characterization of genes in SLG/SRK region of S(9) haplotype of Brassica campestris (syn. rapa). Genetics, 153(1), 391–400. 10.1093/genetics/153.1.391.

Takayama, S., & Isogai, A. (2005). Self-incompatibility in plants. Annu Rev Plant Biol. 56, 467–489. 10.1093/10.1146/annurev.arplant.56.032604.144249.

Tamura, K., Peterson, D., Peterson, N., Stecher, G., Nei, M., & Kumar, S. (2011). MEGA5: molecular evolutionary genetics analysis using maximum likelihood, evolutionary distance, and maximum parsimony methods. Mol Biol Evol. 28(10), 2731–2739. 10.1093/molbev/msr121.

Tang, H., Krishnakumar, V., & Li, J., & Zhang, X. (2015). jcvi: JCVI utility libraries: Zenodo. 10.5281/zenodo.31631. 10.5281/ZENODO.31631.

Velasco R, Zharkikh A, Affourtit J, Dhingra A, Cestaro A, Kalyanaraman A, Fontana P, Bhatnagar SK, Troggio M, Pruss D, et al. (2010). The genome of the domesticated apple (*Malus × domestica Borkh*.). Nat Genet. 42: 833–839. 10.1038/ng.654.

Vieira, Fonseca, N. A., & Vieira, C. P. (2008). An S-RNase-based gametophytic self-Incompatibility system evolved only once in eudicots. J Mol Evol. 67(2), 179–190. 10.1007/s00239-008-9137-x.

Vieira, Pimenta, J., Gomes, A., Laia, J., Rocha, S., Heitzler, P., & Vieira, C. P. (2021). The identification of the Rosa S-locus and implications on the evolution of the Rosaceae gametophytic self-incompatibility systems. Sci Rep. 11(1), 3710. 10.1038/s41598-021-83243-8.

Wang, Y., Tang, H., Debarry, J. D., Tan, X., Li, J., Wang, X., Lee, T. H., Jin, H., Marler, B., Guo, H., Kissinger, J. C., & Paterson, A. H. (2012). MCScanX: a toolkit for detection and evolutionary analysis of gene synteny and collinearity. Nucleic Acids Res, 40(7), e49. 10.1093/nar/gkr1293.

Wheeler, M. J., de Graaf, B. H., Hadjiosif, N., Perry, R. M., Poulter, N. S., Osman, K., Vatovec, S., Harper, A., Franklin, F. C., & Franklin-Tong, V. E. (2009). Identification of the pollen self-incompatibility determinant in Papaver rhoeas. Nature, 459(7249), 992–995. 10.1038/nature08027.

Wu, S., Han, B., & Jiao, Y. (2020). Genetic Contribution of Paleopolyploidy to Adaptive Evolution in Angiosperms. Mol Plant, 13(1), 59–71. 10.1016/j.molp.2019.10.012.

Xiang, Y., Huang, C. H., Hu, Y., Wen, J., Li, S., Yi, T., Chen, H., Xiang, J., & Ma, H. (2017). Evolution of Rosaceae fruit types based on nuclear phylogeny in the context of geological times and genome duplication. Mol Biol Evol. 34(2), 262–281. 10.1093/molbev/msw242.

Zhao, H., Zhang, Y., Zhang, H., Song, Y., Zhao, F., Zhang, Y., Zhu, S., Zhang, H., Zhou, Z., Guo, H., Li, M., Li, J., Gao, Q., Han, Q., Huang, H., Copsey, L., Li, Q., Chen, H., Coen, E., Zhang, Y., & Xue, Y. (2022). Origin, loss, and regain of self-incompatibility in angiosperms. Plant Cell. 34(1), 579–596. 10.1093/plcell/koab266.

Zhao, T., Holmer, R., de Bruijn, S., Angenent, G. C., van den Burg, H. A., & Schranz, M. E. (2017). Phylogenomic synteny network analysis of MADS-Box transcription factor genes reveals lineage-specific transpositions, ancient tandem duplications, and deep positional conservation. Plant Cell. 29(6), 1278–1292. 10.1105/tpc.17.00312.

Zhao, T., & Schranz, M. E. (2017). Network approaches for plant phylogenomic synteny analysis. Curr Opin Plant Biol. 36, 129–134. 10.1016/j.pbi.2017.03.001.

Zhao, T., & Schranz, M. E. (2019). Network-based microsynteny analysis identifies major differences and genomic outliers in mammalian and angiosperm genomes. Proc Natl Acad Sci U S A. 116(6), 2165–2174. 10.1073/pnas.1801757116.

Zhu, S., Zhang, Y., Copsy, L., Han, Q., Zheng, D., Coen, E., & Xue, Y. (2023). The snapdragon genomes reveal the evolutionary dynamics of the S-Locus supergene. Mol Biol Evol. 40(4). 10.1093/molbev/msad080.

